# High immigration rates critical for establishing emigration-driven diversity in microbial communities

**DOI:** 10.1101/2023.02.24.529875

**Authors:** Xiaoli Chen, Miaoxiao Wang, Laipeng Luo, Liyun An, Xiaonan Liu, Yuan Fang, Ting Huang, Yong Nie, Xiao-Lei Wu

**Affiliations:** College of Engineering, Peking University, Beijing 100871, China; Institute of Ocean Research, Peking University, Beijing 100871, China; Microbial Systems Ecology Group, Institute of Biogeochemistry and Pollutant Dynamics, Department of Environmental Systems Sciences, ETH Zurich, 8006, Zurich, Switzerland; Department of Environmental Microbiology, Eawag: Swiss Federal Institute of Aquatic Sciences, 8600, Duebendorf, Switzerland; College of Architecture and Environment, Sichuan University, Chengdu, 610000, China; School of Resource and Environmental Engineering, Hefei University of Technology, Hefei 230000, China; Institute of Ecology, Peking University, Beijing 100871, China

**Author notes:** Corresponding author: Research Scientist, College of Engineering, Peking University. Corresponding author: Professor, College of Engineering, Peking University.

**Keywords:** Emigration and immigration, Microbial diversity, Generalized Lotka-Volterra model, Artificial microbial consortium.

## Abstract

Unraveling the mechanisms governing the diversity of ecological communities is a central goal in ecology. While microbial dispersal (including the emigration and immigration process) constitutes an important ecological process, the coupling effects of dispersal and microbial competition in microbial diversity are poorly understood. Here, we investigated how microbial dispersal affects the diversity of microbial communities in the presence of inter-species competition, using a generalized Lotka-Volterra model in combination with experimental investigations. Our model shows that emigration reduces the diversity induced by immigration at low immigration rates. We surprisingly find that it increases the diversity of the community when the immigration rates cross a defined threshold, which we identified as *I_neutral_*. We also found that at high immigration rates, emigration weakens the relative abundance of fast-growing species, and thus enhances the mass effect and increases the diversity. We experimentally confirmed this finding using cocultures of 20 bacterial strains isolated from the soil. Our model further showed that *I_neutral_* exists over a wide range of species pool sizes, growth rates, and interspecies interactions, and decreases with the increasing of species pool size, growth rate, and interspecies interaction. Our work deepens the understanding of the effects of dispersal on diversity of natural communities.

## Introduction

Microorganisms serve essential roles in an enormous range of biological processes, from global biogeochemical cycles to the regulation of human health^1–6^. The maintenance of microbial diversity is crucial to the sustainability of their ecological functions and services^5–7^. However, according to the principle of competitive exclusion, few species can coexist in an unstructured environment with limited resources, which is inconsistent with the observed diversity in natural ecosystems^8^. This contradiction is often referred to as “the paradox of the plankton”^9^. How microbial diversity is maintained remains one of the unanswered fundamental questions in ecology.

Previous studies suggested that both deterministic processes (including species traits, interspecies interactions, and environmental conditions)^10, 11^ and stochastic processes (including drift, dispersal, extinction, and speciation)^12, 13^ govern community assembly, and determine species diversity of ecological communities^14^. However, despite recognizing that ecological processes beyond deterministic factors are important for community assembly, a mechanistic understanding of how stochastic processes shape the diversity of ecological communities remains absent. As a key stochastic process within the assembly of microbial communities, dispersal critically affects the composition and functioning of microbial communities^15, 16^. In general, the effects of dispersal to a local community are understood to involve two fundamental processes: (1) immigration into and (2) emigration out of a community^17^. Immigration of species from other communities creates mass effects, which is defined as the flow of individuals from areas of high success to unfavorable areas^18^. Thus, mass effect may rescue the species from extinction in communities where they are weak competitors^19^ and always increases microbial diversity^20^. In contrast, emigration drives the loss of local populations in closed communities, a process that greatly diminishes microbial diversity^21^. However, little is known about how precisely immigration and emigration affect microbial diversity in a collective manner^22^.

The relationship between dispersal^23^ (including emigration^24^ and immigration^25^) and microbial diversity in natural communities has previously been investigated. The results showed that some species exhibit a tradeoff between growth rate and competitive ability in the biosystem. High emigration favors the species with high growth rates and low competitive abilities, whereas low emigration favors the species with low growth rates and high competitive abilities^24^. However, if species do not exhibit a trade-off between growth rate and competitive ability, and the immigration process is introduced into the community succession, the precise effects of these both dispersal processes on microbial diversity remain to be quantified. The dispersal including immigration and emigration has been evaluated for synthetic bacterial community^23^, but the process of immigration and emigration is coupled. In this study, we decoupled the immigration and emigration process to explore how immigration and emigration individually and jointly affect the diversity of the communities.

Here, we combined a generalized Lotka-Volterra model with a synthetic microbial consortium, to investigate how emigration and immigration in combination regulate the species diversity of a community. We created a synthetic microbial community composed of 20 strains (from 12 genera and 20 species) to emulate an ecological biosystem. To manipulate the dispersal strength (the number of cells dispersal into/out of a community per time, referred to as immigration rate and emigration rate, respectively), coculture experiments were subject to daily growth/dilution cycles across a series of dilution factors and mass resupplies. We find that at low immigration rates, emigration decreases the Shannon diversity of microbial communities; while at high immigration rates, emigration increases the Shannon diversity of microbial communities. Our results highlight that the relative abundance of fast-growing species plays a key role in the regulation of microbial diversity.

## Results

### The simulation predicts that at high immigration rates, emigration increases Shannon diversity of microbial communities

We focus on the regime where the diversity of the communities is determined by interspecies competition and dispersal (including immigration and emigration). To probe the effects of emigration and immigration on species diversity, we constructed a generalized Lotka-Volterra model that incorporates the emigration term δ*N_i_* and immigration term *imm*^24, 26, 27^, as follows:

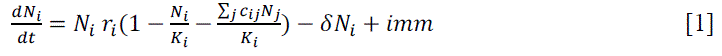

where *N_i_* is the density of species *i*; *r_i_* is the maximum growth rate of species *i*; *K_i_* is the carrying capacity of species *i*; the interaction coefficient *c_ij_* is defined as the effect of species *j* on species *i*. The simulations were initialized with a species pool containing *S* species. To match the following experimental expression, we defined the dilution factor as 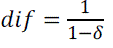, which represents the magnitude of the emigration rate^24^. To study the effect on community succession of the emigration and immigration rates, we finished the model by varying the degree of dilution factor *dif* and immigration rate *imm*. To guarantee the communities reach steady states, we set the time to the simulation time to 3×10^3^ for each simulation (Fig. S3). Microbial diversity was assessed by measuring the Shannon index of the community after simulation^28^.

We first created an ecosystem composed of 10 species. We found that when no immigration occurred, the Shannon index was negatively associated with dilution factors (Fig. 1A, Spearman correlation: *R* = −0.847, *P*-value < 0.001). The rank-abundance curves for the community with high dilution factors were characterized by a more left-skewed abundance distribution (Fig. 1B), indicating that increased emigration of microbial communities resulted in the extinction of low-abundant species within the affected community. Without emigration, the Shannon index positively correlated with the immigration rates (Fig. 1C, Spearman correlation: *R* = −0.781, *P*-value < 0.001). We found that in communities characterized by higher immigration rates, the gradient of the curves flattened (Fig. 1D). This result points to a higher evenness since the relative abundance of different species was more similar in the community. Our model results indicated that immigration increased microbial diversity by rescuing species from local competitive exclusion, which could be explained by mass effect.

**Fig. 1.**
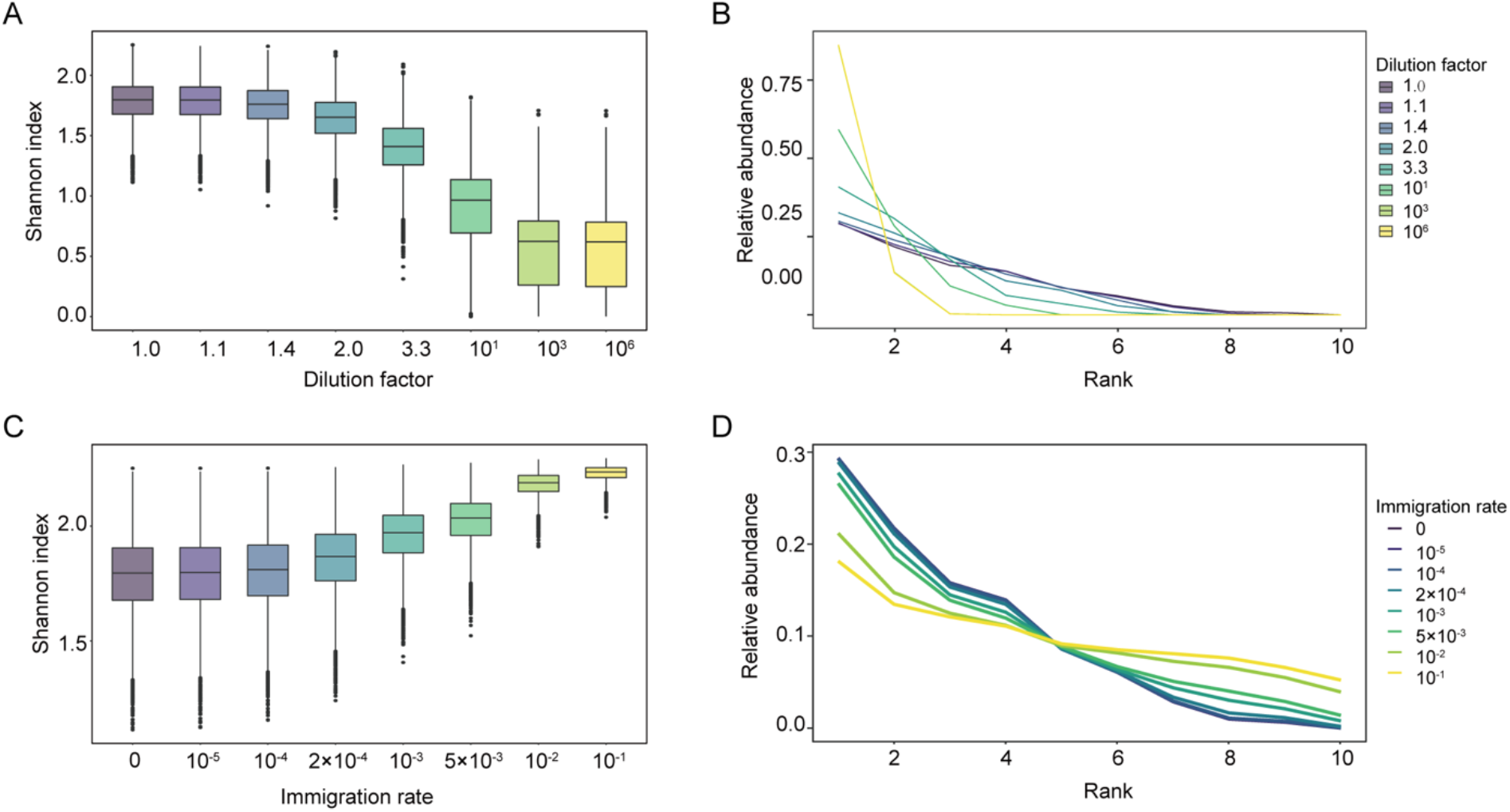
The Lotka-Volterra model predicts that emigration and immigration had opposite effects on the Shannon diversity of microbial communities. The values of δ were set to 0, 0.1, 0.3, 0.5, 0.7, 0.9, 0.999, and 0.999999. The corresponding values of *dif* were 1.0, 1.1, 1.4, 2.0, 3.3, 10^1^, 10^3^, and 10^6^ in (A) and (B). (A) The Shannon index of microbial communities negatively correlates with the dilution factors. (B) The effect of emigration on rank-abundance curves for the communities. The X-axis of the curve represents species rank and Y-axis represents the relative abundance of species in a community. (C) The Shannon index of communities positively correlates with immigration rates. (D) The rank-abundance distributions of the communities across a range of immigration rates. Immigration makes the species’ abundance distributions even. Figures were drawn using 10^3^ independent simulations. For each simulation, we create an ecosystem composed of 10 species. Parameter values used in this simulation: interactions strengths (*c*_$(_), independently drawn from normal distribution with a mean of 0.5 and a standard deviation of 0.1; carrying capacities (*K_i_*), independently drawn from normal distribution with a mean of 2 and a standard deviation of 0.2; growth rates (*r_i_*), independently drawn from normal distribution with a mean of 1 and a standard deviation of 0.1.

Next, we investigate the effects of both immigration and emigration on the diversity of microbial communities. To avoid the situation that immigration and emigration are both very strong so that the term in parentheses 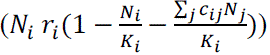 in Equation [1] are ignored (Fig. S4), the immigration rates were set to ≤ 10^*+^, lower than species’ growth rates. Notably, when we allowed both emigration and immigration to occur in a microbial community, we observed that increasing the dilution factors resulted in a decrease in Shannon diversity at lower immigration rates (≤ 3 × 10^*,^). In contrast, increasing the dilution factors resulted in an increase in Shannon diversity at higher immigration rates (≥ 8 × 10^*,^) (Fig. 2A). As shown in Fig. 2B, at high immigration rates (5 × 10^*,^ and 10^*-^), the rank-abundance curves for the community with a high dilution factor (10.) displayed a less steep decline than that with a low dilution factor (10^+^), indicating higher Shannon diversity for the community with a high dilution factor (10.). A value of immigration rate (simplified as *I_neutral_*) was calculated, which represents the Shannon indexes of communities characterized by high dilution factors which are equal to those of communities characterized by low dilution factors. When the immigration rate is over the value of *I_neutral_*, emigration increases the Shannon diversity of the community. To obtain the value of *I_neutral_*, we first calculated the slope at each immigration rate by linear fitting of dilution factors and Shannon indexes. Second, we obtained the curve of the slope changing with the immigration rates by nonlinear fitting of the immigration rates and these slopes (See Mathods). The immigration rate on this curve with a slope of zero represents the *I_neutral_* (Fig. 2C). When we applied this method to calculate the *I_neutral_*, we obtained the logarithm of *I_neutral_* of –2.25 in these simulations. Together, these results demonstrated that emigration increased the Shannon diversity of the community when the immigration rate was above *I_neutral_*.

**Fig. 2.**
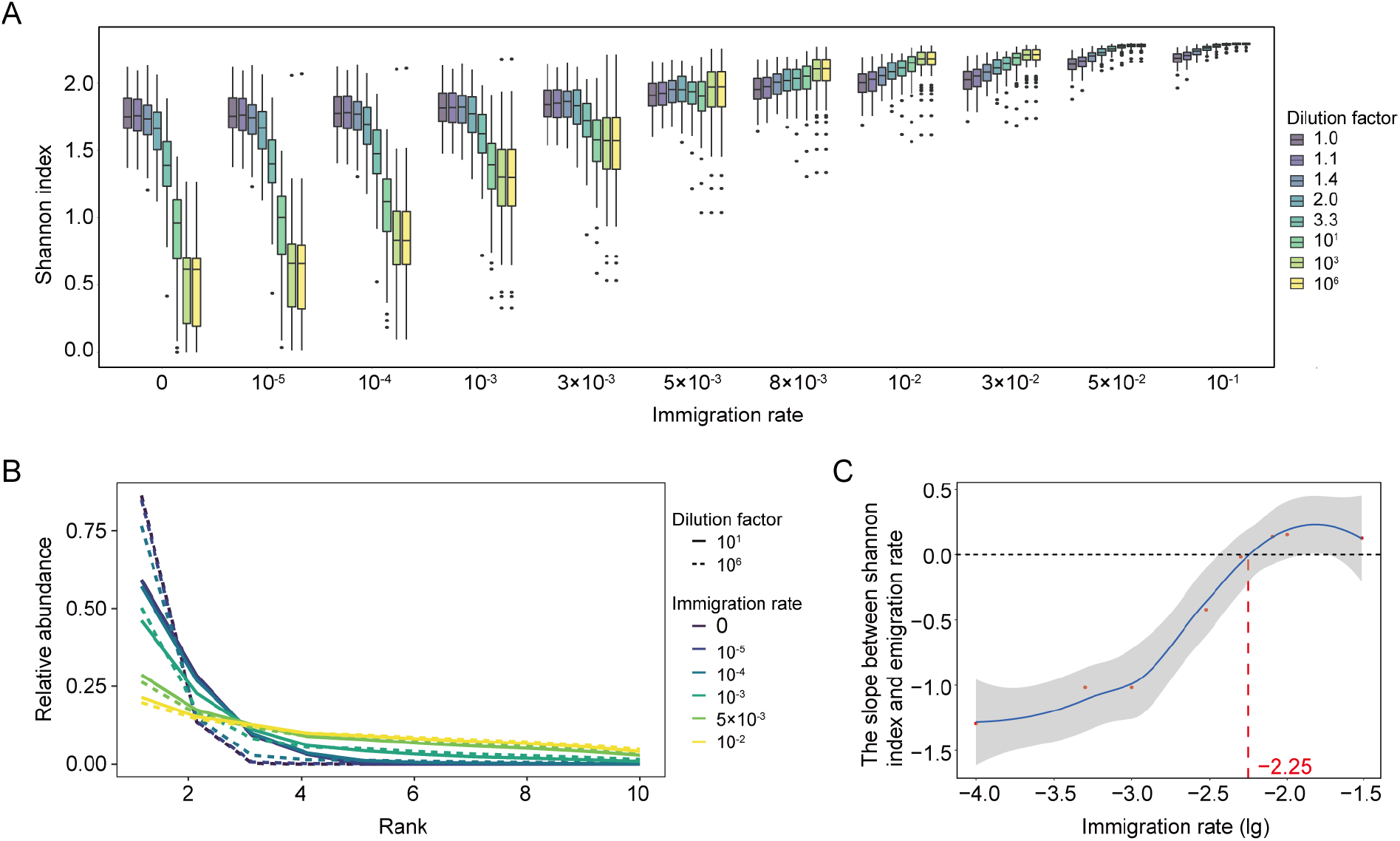
The Lotka-Volterra model predicts that an increasing immigration rate forces the diversity of the high-emigration community to exceed the diversity of the low-emigration community. (A) At lower immigration rates (≤3×10^-3^), Shannon index decreased with increasing dilution factors. At higher immigration rates (≥8×10^-3^), Shannon index increased with increasing dilution factors. (B) Effects of both dilution factors and immigration rates on rank-abundance curves for the communities. (C) The X-axis represents the logarithm of the immigration rate. The Y-axis represents the slope of the linear fit between the Shannon index and the dilution factors at each immigration rate. The value of *I_neutral_* (2.25) represents the immigration rate at the slope of zero fitted between the Shannon index and dilution factor. Figures were drawn using 10^3^ independent simulations. Parameter values used in Fig. 2 are identical to those used in Fig. 1.

## Computational simulations reveal how dispersal affects species diversity

To better understand why emigration increases the Shannon diversity of the community when the immigration rate is above *I_neutral_*, we analyzed the role of growth characteristics of species (i.e., growth rate) in community succession. We fitted the relationship between the growth rate of each species in the community and its abundance across a range of emigration and immigration rates. We found that under all simulated immigration and emigration conditions, the slopes obtained by linear fitting of growth rate and relative abundance of species in the community (Fig. 3A) were greater than zero, manifested as significant positive correlations between the growth rate and abundance of species (Fig. 3B, linear regression, *R*^2^ ≥ 0.37, *P*-value < 0.001). The values of slope reflect the abundance advantages as a result of high growth rates of the species in the community. As shown in Fig. 3B, the slopes decreased with the increasing immigration rates, both under high and low dilution factors, indicating that immigration weakened the advantages of species as a result of high growth rates in the community. Notably, the slopes with a high dilution factor of 10. were larger than that with a low dilution factor of 10^+^ when the immigration rates were lower than ≤ 10^*,^, which indicated that emigration favored fast-growing species at low immigration rates. In contrast, we found that when the immigration rates were over 5 × 10^*,^, the slopes with a high dilution factor of 10. were lower than that with a low dilution factor of 10^+^, indicating that emigration reduced the advantage of relative abundance of fast-growing species at high immigration rates. This result indicated that the advantages of the fast-growing species were diminished by high emigration at high immigration rates, which resulted in increased Shannon diversity through emigration at high immigration rates (Fig. 2A).

**Fig. 3.**
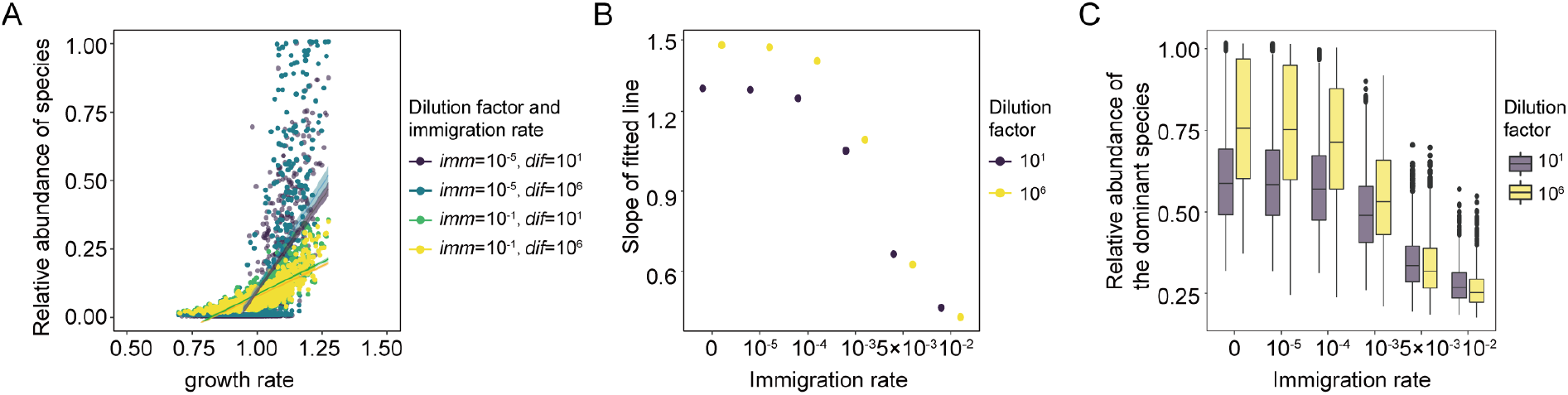
Results of the 10-species model showing that the dispersal process influences advantages as a result of species’ growth rate, and further influences the relative abundance of dominant species featuring high growth rates. (A) Linear regressions between the relative abundance of species and growth rates of species in the community at different dilution factors and immigration rates. (B) Effects of both emigration and immigration on the slopes of linear regressions between the relative abundance of species and growth rates of species in the community. (C) Effects of both emigration and immigration on the relative abundance of dominant species in the communities. Parameter values used in Fig. 3 are identical to those used in Fig. 1.

We next investigated how the advantage as a result of the high growth rate of species affected the relative abundance of species in a community. To this end, we analyzed the changes in the relative abundance of dominant species (the most abundant species in the community)^29^, which represents the relative abundance of low-ranking species in of rank-abundance curve. As shown in Fig. S5, the dominant species exhibited faster growth rates compared to the other species of the community assessed. We found that the relative abundance of the dominant species in the community decreased with increasing immigration rates regardless of dilution factors (Fig. 3C), which might be caused by a decline in the advantage of the fast-growing species (Fig. 3B). At low immigration rates (≤ 10^*,^), the relative abundance of dominant species increased with the increasing dilution factors (Fig. 3C). In contrast, at high immigration rates, the Shannon index of the community decreased with the increasing dilution factors (Fig. 2A). As the immigration rate increased to 5 × 10^*,^, the relative abundance of dominant species in a community with a high dilution factor of 10. was lower than that with a low dilution factor of 10^+^ (Fig. 3C), resulting in increasing the Shannon diversity of the high-emigration community more than that of the low-emigration community at high immigration rates (Fig. 2A). Together, our simulation results showed that at high immigration rates, emigration increased the Shannon diversity of communities, accompanied by a lower relative abundance of dominant species featuring high growth rates.

## Confirmation of our Model observations by testing a synthetic consortium

## 1. Monotonic effects of single dispersal factors on microbial diversity

To experimentally test whether emigration would enhance the Shannon diversity of microbial communities at high immigration levels, we assembled a synthetic consortium containing 20 bacterial isolates from a soil sample taken near Weiming Lake at Peking University (Fig. S1)^30^. To mimic its assembly process, we passaged the consortium across six daily growth/dilution cycles. To regulate the levels of emigration, we changed dilution factor during the passages, with a higher dilution factor mimicking a higher emigration rate (Fig. 4A)^24^. Immigration with different biomass was imposed by adding the 20-strain pool to the local community at the beginning of each growth cycle (Fig. 4B-C).

**Fig. 4.**
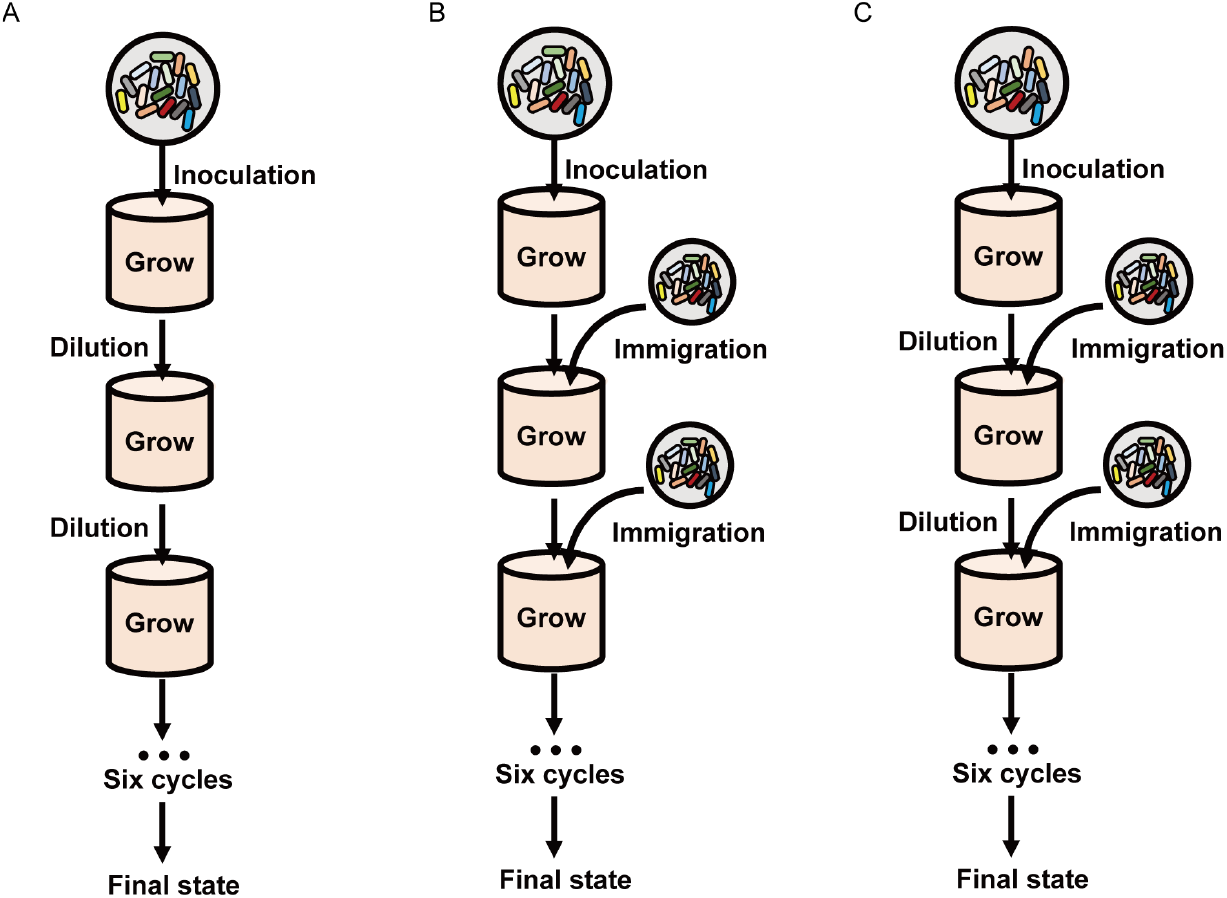
Schematic of the growth-dilution-immigration scheme employed in co-cultural experiments. The regional pool was composed of 20 species at equal ratios. The microbial communities were grown until a stable equilibrium was reached. (A) Diagram showing how the communities were diluted over a range of dilution factors during succession. The dilution factor determined the fraction of cells discarded, and thus the emigration rates. (B) An immigration community containing all species at equal ratios at multiple immigration rates was added to the cultures during succession. (C) The communities were grown in the presence of both dilution and immigration. Dilution factors and immigration rates were changed over several orders of magnitude.

We first used our system to examine how emigration or immigration individually affects species diversity within the community. When we excluded the immigration process, we found that the Shannon index of the consortia negatively correlated with the dilution factors (Fig. 5A, Spearman correlation: *R* = −0.831, *P*-value < 0.001). In the rank-abundance curves for the community with different dilution factors, the ranks corresponding to abundance greater than 0 with high dilution factors (10^/^ and 10.) were lower than that with low dilution factors (10^+^ and 10^-^) (Fig. 5B), suggesting that emigration significantly decreased the Shannon diversity of microbial communities by species loss. To determine how immigration from a source pool affected species diversity, we eliminated “bottleneck effects” by setting the dilution factor to a low level (10^1^) to eliminate the, before adding the 20-strain pool to the local community under different immigration rates. Our results showed that the Shannon index positively correlated with immigration rates (Fig. 5C, Spearman correlation: *R* = 0.784, *P*-value < 0.001). The steepness of the abundance rank curves decreased with the increase in immigration rates. In particular, when the immigration rate increased to 10^1^, the species richness in the community was significantly higher than that without immigration (Fig. 5D and Fig. S6), suggesting that immigration improved the diversity of the communities through species recolonization. Together, the fundamental effects of a single dispersal process on microbial diversity in our experimental system were in agreement with the effects observed in our model (Fig. 1). Based on these findings, we next investigated the combined effects of the two dispersal processes, including emigration and immigration on the diversity of microbial communities.

**Fig. 5.**
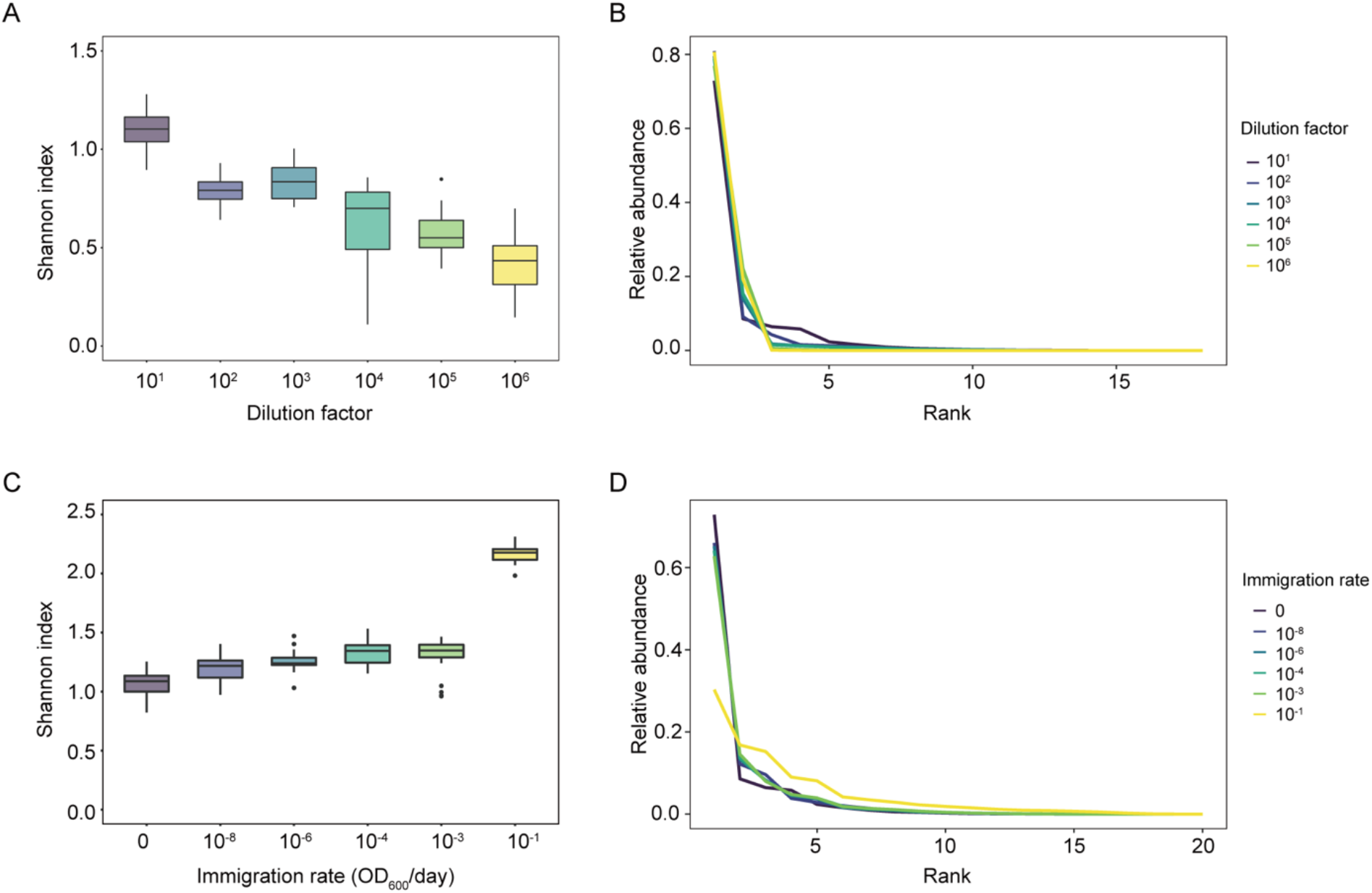
Impact of the emigration alone and immigration alone on the diversity of bacterial communities and the rank-abundance curves for the communities in the experimental system. (A) Emigration decreases the Shannon diversity of bacterial communities in the absence of immigration. (B) Rank-abundance plot of the communities under a series of dilution factors. (C) Immigration increases the Shannon diversity of bacterial communities. The consortium was cultured through daily growth-dilution cycles under the dilution factor of 10. (D) The effect of immigration on rank-abundance curves for the communities.

## 2. Combined effects of immigration and emigration on the diversity of bacterial communities

To assess the combined effects of immigration and emigration in combination on the species diversity of the community, we set up a synthetic consortium consisting of 20 species as described above and passaged the co-cultures through daily growth-dilution cycles under two dilution factors (10^1^ and 10^6^) and a series of immigration rates. We found that the Shannon index of the microbial community with a lower dilution factor of 10^1^ was higher than that with a higher dilution factor of 10^6^ in the co-cultures without immigration or with low immigration rates (≤ 10^*.^). However, when we increased the immigration rates to 10*^0^, the Shannon diversity of the community with a high dilution factor of 10^6^ was higher than that with a low dilution factor of 10^1^ (Fig. 6A, Wilcoxon test, *P*-value < 0.001). In agreement with the simulation results (Fig. 2B), under the condition of high immigration rates (≥ 10^*,^), the rank-abundance curves for the high-emigration community were less steep than those for the low-emigration community (Fig. 6B). In this regime, the relative abundance of low-ranking species at the head of the curve in low-emigration communities was lower than those in high-emigration communities. In addition, the relative abundance of rare species at the tail of the curve in high-emigration communities was higher than those in low-emigration communities. These abundance changes of dominant species and rare species under high immigration resulted in higher Shannon diversity of high-emigration communities. The experimental results confirmed the presence of *I_neutral_* over which emigration favored the diversity of microbial communities. Among the 20 strains in the experimental biosystem, the growth rate of Ka was significantly higher than that of the other 19 strains (Fig. S7). Ka dominated the microbial community under the condition of low immigration rates (≤ 10^*.^) (Fig. 6C and Fig. S6), which indicates that a high growth rate resulted in an advantage in the relative abundance of the species in the community. To investigate the relationship between the relative abundance of fast-growing species and the Shannon diversity in the community, the abundance of Ka was calculated. At low immigration rates (≤ 10^*.^), the relative abundance of fast-growing species (i.e., Ka in the experimental biosystem) remained at a high level in the microbial community with a dilution factor of 10^6^ (Fig. 6C), while the Shannon index of the microbial community was low. In comparison, at high immigration rates (≥ 10*^0^), the relative abundance of the fast-growing species in the microbial community with a dilution factor of 10^6^ was lower than that with a dilution factor of 10^1^, resulting in the Shannon index at a high dilution factor of 10^6^ higher than that at a low dilution factor of 10^1^. This result suggested that high immigration rates decreased the relative abundance of fast-growing species in a high dilution factor more greatly than that in a low dilution factor, and the diversity of high emigration community was greater than that in low emigration community. Overall, our experimental results confirmed our model observations, which predicted that under the high immigration rates, the abundance of fast-growing bacteria decreased with increasing dilution factors, resulting in higher Shannon diversity in the high-emigration community than in the low-emigration community.

**Fig. 6.**
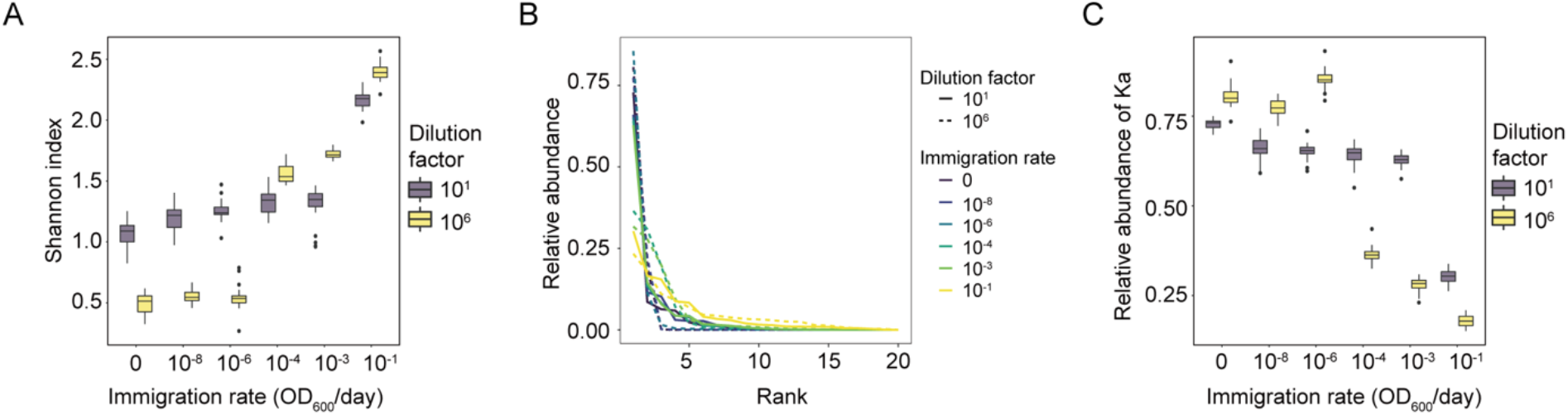
Experimental results showing effects of the dilution and immigration rates on the diversity of bacterial communities and the relative abundance of the fast-growing species (Ka). (A) Effects of both dilution and immigration rates on the Shannon diversity of bacterial communities. (B) The effect of both emigration and immigration on species abundance distributions of the microbial communities. (C) Relative abundance of the fast-growing species under different dilution factors and across a range of immigration rates.

## Extended computational simulations reveal that *I_neutral_* exists cover a wide range of parameter values

As our model identified an *I_neural_* at which emigration rates did not affect the ecosystem diversity in one parameter set, we tested whether the observed *I_neutral_* existed in the communities by varying key parameters, including the number of species, interaction strengths, and growth rates. First, we employed the Lotka-Volterra model with different species pool sizes. As shown in Fig. 7A, the *I_neutral_* decreased with increasing species pool size. Next, to identify how interaction strength *c_ij_* and growth rate *r_i_* of species *i* determined the *I_neutral_* associated with the assembly of a given community, we used a 10-species biosystem. The model results showed that *I_neutral_* negatively correlated with *c_ij_* while fixing the growth rate of species (Fig. 7B). We also found that *I_neutral_* significantly changed whether the values for interaction strength were high or low. In contrast, *I_neutral_* had a slight downward trend at intermediate levels of interaction strength (*c_ij_* = 0.3∼0.8), suggesting that species’ interaction strength over a wide range had a negligible effect on *I_neutral_*. When we changed growth rates of species, we found that the increasing growth rates of species for a given interaction strength resulted in a lower *I_neutral_*. In addition, we found that as the growth rate increased, *I_neutral_* steadily decreased (Fig. 7C). Our results indicated that *I_neutral_* exists over a wide range of ecological parameters (i.e., varying species pool size, interaction strength, and growth rate), despite the fact that several key ecological parameters (species pool size, interaction strength, and growth rate) of a microbial community can regulate the effects of dispersal on biodiversity.

**Fig. 7.**
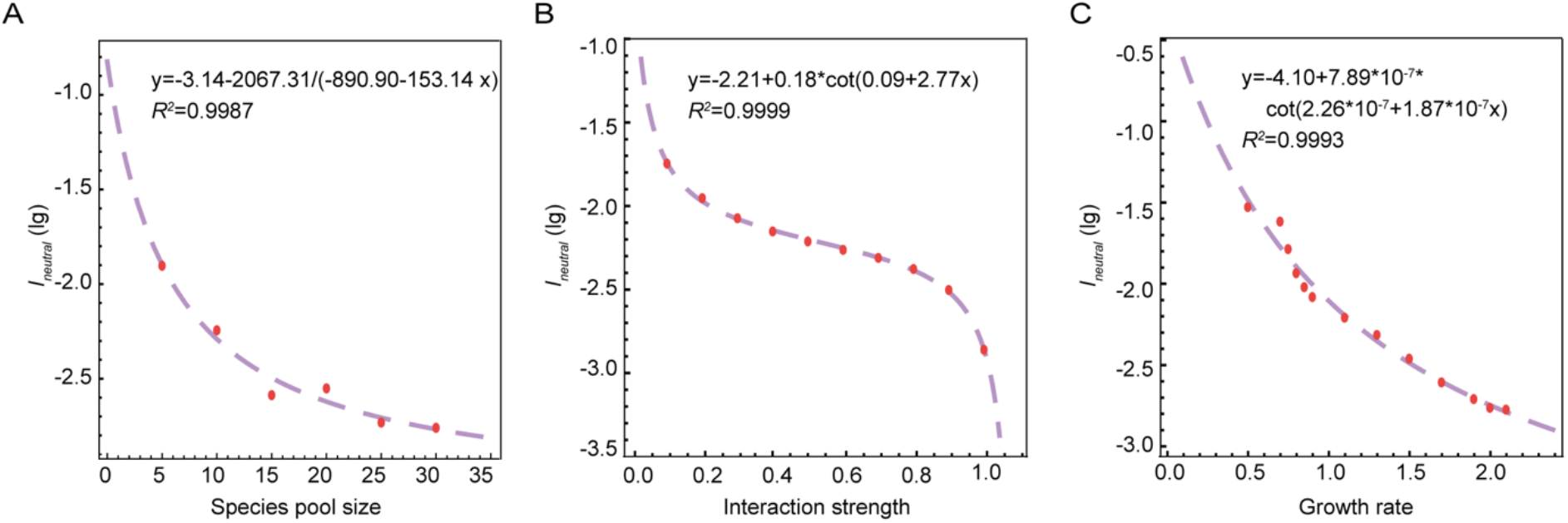
Species pool size, interaction strength, and growth rate determined *I_neutral_*. *I_neutral_* negatively correlated with the parameters of species pool size (A), interaction strength (B), and growth rate (C). (A) Effects of species pool size on Shannon diversity while fixing interaction strength (<*c*_$(_>=0.5, std (*c*_$(_)=0.1) and growth rate (<*r_i_*>=1, std (*r_i_*)=0.1). (B) Effects of interaction strength on Shannon diversity while fixing the species pool size (i.e, 10 species) and growth rate (<*r_i_*>=1, std (*r_i_*)=0.1). (C) Effects of growth rate on Shannon diversity while fixing the species pool size (i.e, 10 species) and interaction strength (<*c*_$(_>=0.5, std (*c*_$(_)=0.1). The dots in the plots represent the mean value over 1000 simulations.

## Discussion

Here, we used both model and experiment to show that the loss of species with increasing dilution factors led to a decrease in the Shannon diversity of an ecosystem in the absence of immigration. However, the effect of emigration levels on microbial diversity shifted from decreasing to increasing with increasing immigration. At high immigration levels, emigration weakened the relative abundance of fast-growing species, thus improving the Shannon diversity of the community. Using the generalized Lotka-Volterra model, we further quantified the impact of key parameters (i.e., species pool size, interaction strength, and growth rate) on *I_neutral_*, which represents the Shannon indexes of communities characterized by high dilution factors which are equal to those of communities characterized by low dilution factors.

Empirical and theoretical studies of biosystems have previously demonstrated that the size of communities is negatively correlated with ecological drift^31–33^. It is well-established that low-abundance microorganisms are more vulnerable to the effects of ecological drift, as slight negative changes in their abundance are likely to result in their extinction^34, 35^. In our system, ecological drift plays a stronger role when low-abundance species in communities go randomly extinct with increasing dilution factors, driving microbial diversity with high dilution factors smaller than those with low dilution factors. Our model predicts that fast growers have the advantage of being the majority in the community regardless of emigration levels, which is inconsistent with previous observations^24^. A previous study showed that at high emigration levels, slow growers with high competition coefficients dominated the community^24^. In our model, however, the competition coefficients of interaction forces of fast growers and slow growers are similar. As a result, the slow growers do not have the advantage at high levels of emigration out of the community.

Our findings are consistent with previous studies. These studies claimed that immigration allows bacterial populations to constantly colonize the localities within the local community, causing the rescue effects, and thus substantially mitigating diversity loss. In previous studies, immigration was a major contributor to buffering against extinction, primarily by recolonization. In those studies, as immigration rates increase, an organism with a competitive disadvantage can be sustained in the microbial system. Conversely, the species with a competitive advantage is more able to dominate the local community^23, 36–38^.

Our study examined the role of both emigration and immigration simultaneously. We found that emigration decreased the diversity of the microbial communities at low immigration rates, but increased the diversity of the microbial communities at high immigration rates, different from previous studies focusing on just one single factor^39, 40^. Our model and experimental results demonstrate that with the increase in immigration rates, the diversity of the high-emigration community was higher than that of the low-emigration community. High rates of propagule input coupled with small local communities provide a mechanism whereby inferior competitors enter resident communities, thus increasing microbial diversity^32^. This enabled us to observe that under high immigration rates, the diversity of high-emigration communities was higher than that of low-emigration communities. We found that when a new community was added, at low immigration rates, the small input of external communities cannot homogenize the resident community. However, our results clearly show that at high immigration rates, the resident community with a high dilution factor is overwhelmed by mass effects, resulting in the existence of *I_neutral_*. Specifically, the main reason responsible for the existence of *I_neutral_* is that high dilution factors correspond to a small inoculation community size, which is more vulnerable to mass effects^31^. As a consequence of mass effects, the relative abundance of superior native competitors declines, and more inferior competitors colonize, resulting in high Shannon diversity of the microbial community.

Previous studies revealed that the steady-state diversity was strongly dependent on the dynamics of the microbial community on both the species pool size and interaction strength^41^. In our study, in addition to these two parameters, we also determined the effects of dispersal on community diversity when species in communities were set at different growth rates. *I_neutral_* is present over a wide range of these parameters. *I_neutral_* decreased with increasing these parameters. However, we found here that *I_neutral_* differs as a function of species pool size, interaction strengths, and growth rates. Specifically, *I_neutral_* changes slightly with the interaction strength over a wide range of interaction strengths (*c_ij_* = 0.3∼0.8), while *I_neutral_* changes largely with the growth rate over a wide range of growth rates (*r_i_* = 0.5∼1.5). One reason for the observed difference in *I_neutral_* as a function of interaction strengths or growth rate may result from the fact that the dominant species show an advantage in growth rate rather than an advantage in interaction strength within the community. Our results suggest that it is possible to quantitatively predict the effect of dispersal on microbial diversity by several key features, such as the species pool size, the statistics of interaction strengths and growth rates.

Although we have found that the dispersal process played a crucial role in controlling microbial diversity, several limitations in our system must be addressed in future work. Our study is based on simple synthetic communities composed of 20 species, which cannot reproduce the complexity of natural systems. For example, our experimental system only includes cooperation and competition interactions. In particular, the Lotka-Volterra model only contains competition interactions, while other interactions such as predation and symbiosis are not included. In addition, in the presence of immigration, only two dilution factors (10^1^ and 10^6^) were performed in our experiments. To calculate the *I_neutral_* in the experimental systems, it is necessary to do more experiments with more dilution factors. The results of species pool size, growth rate, and interaction strength affecting *I_neutral_* should also be further verified experimentally.

## Conclusion

In summary, we provide new insight into how microbial diversity depends on both emigration and immigration. At high immigration levels, emigration decreases an advantage in the relative abundance of fast growers, resulting in the increased Shannon diversity of the community. We demonstrate that the effect of immigration and emigration on community assembly is not independent, which enlightens us that when we study both immigration and emigration process, we should consider the two process as a whole. We further quantified the coupling effect of immigration and emigration on the Shannon diversity of microbial communities using mathematical model. Our study suggests that emigration accompanied by high immigration levels may be a strategy for natural microbial communities to regain diversity.

## Materials and Methods

### Lotka-Volterra simulations

In agreement with previous reports^24^, our model assumes that the growth of each species within a microbial community is determined by the combination of its intrinsic growth rate (*r_i_*), its carrying capacity (*K_i_*), and the combination of any interactions each species has with other community members (*c_ij_*). Beginning with equal abundances of all species, the dynamics of microbial communities were simulated with different species pool sizes *S* ranged from 5 to 30, interaction matrices, and the maximum growth rate of species *i*. For each simulation, the parameter *c_ij_* were independently drawn from a normal distribution with a mean ranging from 0.1 to 0.9 and a standard deviation of 0.1^26^, and a set of *r_i_*were created from a normal distribution with a mean ranged from 0.2 to 2 and a standard deviation of 0.1.

## Calculation of Shannon index

We determined the diversity of each microbial community by calculating its Shannon index *H*:

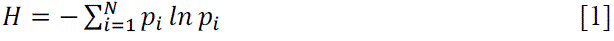

where *p_i_* is the frequency of the *ith* bacterium, *N* is the total number of bacteria in a community.

## Calculation of the value of immigration rate (***I_neutral_***) at which the effects of emigration levels on microbial diversity switch from a decrease to an increase

Linear fitting was performed between Shannon indexes and dilution factors at each immigration rate and the slope at each immigration rate was obtained. Next, locally weighted regression was performed between slopes and immigration rates using the *loess* method of *R*. The immigration rate at slope 0 was defined as the *I_neutral_* for the specific simulation condition. By fitting species pool size, interaction coefficient, or the growth rate of species *i* and *I_neutral_*, the function relation between them was obtained.

## Strain isolation and identification

Bacterial strains were isolated from one identical location of soil environment near Weiming Lake at Peking University. Isolated colonies were cultured in Luria Bertani (LB) medium (0.5% yeast extract, 1% tryptone, 1% NaCl) at 30℃ until strains grew to stationary phases. All strains were cultured while shaking at 220 rpm. Strains that had been cultured were stored in 20% glycerol at −80℃ until further use.

The 16S rRNA gene was sequenced via Sanger sequencing using the universal 16S rRNA primers (i.e., 8F and 1492R), yielding assembled sequences of ∼1200 nt in usable length. Species names were assigned using the National Center for Biotechnology Information (NCBI) database based on the type strain with the highest sequence similarity relative to the query strain (Table S1). For phylogenetic analysis, sequences of 16S rRNA gene were aligned using MEGA 7.0.21 and a tree was constructed by the neighbor-joining method.

## Correction for differences in DNA extraction efficiency

Since the selected 20 strains, namely 3 strains of Gram-positive bacteria and 17 strains of Gram-negative bacteria, originated from different genera, we found a bias in DNA extraction due to different lysis efficiencies. Therefore, a reference strain was introduced when DNA was extracted. The DNA extraction efficiency of the test strain was corrected by mixing a certain proportion of the test strain and the reference strain, extracting DNA, and measuring the ratio of the number of DNA amplicons between the test strain and the reference strain. The correction coefficient of extraction efficiency was calculated to facilitate data correction in the following data analysis. The DNA extractions were performed by combining steel ball crushing and magnetic bead purification from Beijing Biotech Biotechnology Co., Ltd. Details about the DNA extraction efficiency correction are described in Supporting Information: S1.1–S1.2. The obtained DNA was used for 16S amplicon sequencing of the V3-V4 region.

Our reference strain was selected to meet both the following two requirements: 1) The reference strain must be easy to obtain and cultivate; 2) The V3V4 region of the 16S rRNA sequence of the reference strain can be distinguished from the 20 strains used in the experiment. Strain Pg, which is the most similar to *Agrobacterium tumefaciens* GV 3101 among the 20 strains, has only 82.97% sequence similarity in the 16S rRNA V3-V4 region. Based on these conditions, *A. tumefaciens* GV 3101 was chosen as the reference strain.

## Emigration and immigration passaging experiments

Frozen stocks of individual species were streaked out on LB agar Petri dishes, grown at 30℃ until strains grew up, 36 h for Cc, Fg, Pk, and Pa; 72 h for Kr; 12 h for all other strains (The detailed description of the strains was shown in Table S1). Before we performed emigration and immigration passaging experiments, single colonies were picked and each species was grown separately in 3 ml LB broth until their stationary phases (ranging from 12 to 24 h for different strains), and then 2% inoculum was transferred to 20 ml LB broth for 24 h. Cells were then collected by centrifuging at 3000 g for 10 minutes and were washed three times using M9 minimal medium (containing M9 salts, 13.52 mg/L FeCl_3_·6H_2_O, 1.98 mg/L MnCl_2_·4H_2_O, 1.36 mg/L ZnCl_2_, 0.48 mg/L CoCl_2_·6H_2_O, 0.34 mg/L CuCl_2_·2H_2_O, 0.48 mg/L NiCl_2_·6H_2_O, 0.48 mg/L Na_2_MoO_4_·2H_2_O, 0.35 mg/L Na_2_O_3_Se, 0.12 mg/L H_3_BO_3_, 0.24 g/L MgSO_4_, 0.01 g/L CaCl_2_) without carbon source. The individual cultures were then adjusted to an OD_600_ = 2 and mixed at equal proportions to obtain pre-cultures. The pre-cultures were diluted in M9 minimal medium (containing 4.5 g/L glucose) with a final OD_600_ of 0.1. During the emigration and immigration passaging experiments, cell cultures were incubated in 400 μL 96-well plates (NEST), with each well containing a 120-μL culture. They were propagated via serial passaging once per day across a broad range of dilutions at 30°C and shaken at 600 rpm (Thermal Shaker, AOSHENG, China). After each passage, the remainder of the cultures were stored in 20% glycerol and frozen at −80℃ for DNA extraction, amplification, and sequencing. These transfers were repeated 24 times for each independent experiment carried out. Community composition was measured at each transfer.

## Measurement of the growth rate

To determine the growth of 20 species in our experiments, we performed monocultures in M9 minimal medium with glucose (4.5 g/L) as the sole carbon source. Cell density (OD_600_) was measured by the plate reader at regular intervals (i.e., 2 h) over the course of 24 h. Maximum growth rates were determined by averaging growth curves of three replicates using the logistic equation^42^:

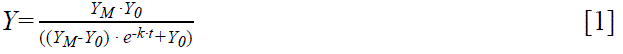

where *Y_0_*is the minimum population biomass; *Y_M_* is the maximum population biomass; and *k* represents the maximum rate of biomass increase (Growth rate; unit, OD_600_ h^-1^).

## DNA amplification and high-throughput sequencing

To reveal the composition of each culture, 72∼96 culture samples were merged into one mixed sample for library construction and sequencing. Barcodes were added to the front of the forward primer for each sample in the library, which were used to split samples in the library later. For the library preparation, amplification of the V3-V4 region of the bacterial 16S rRNA gene was performed to assess the bacterial community using the barcoded primers 338 F: 5′-Barcode+ACTCCTACGGGAGGCAGCA-3′ and 805 R: 5′-GACTACHVGGGTATCTAATCC-3′. After PCR amplification of the extracted DNA, DNA electrophoresis was performed. The brightness of the bands running through electrophoresis reflects the DNA concentration. The concentration of all samples was adjusted to the same by adding ddH_2_O. 72∼96 samples in the same plate were then mixed in equal volumes. The procedure of adjusting samples’ concentration according to band brightness was only to ensure that the samples with low concentrations could receive sufficient sequencing reads. DNA in samples was amplified, purified, and finally sequenced on Illumina HiSeq 2500 platform at Guangdong Magigene Biotechnology, China^43^.

## 16S rRNA gene sequence processing and statistical analyses

The barcodes of the raw sequencing reads were removed using QIIME (v1.9)^44^, generating fastq files with the forward and reverse reads. The sequencing data were then demultiplexed, denoised, and clustered into amplicon sequence variants (ASVs) using the QIIME 2 pipeline (v2020.2)^45^. The ASVs were aligned to the 16S rRNA gene sequence of 21 strains (including 20 strains and a reference strain) to obtain the species composition information of each sample. The α-diversity analysis was performed using R (v3.6.2). Unless indicated otherwise, the number of replicates was 10^3^ for each simulation and 24 for each experiment. For comparative statistics, the *P* values were obtained using an unpaired, two-tailed, Student’s t-test.

## Data availability

The source codes used for our mathematical modelling are available at Github: https://github.com/chenxiaolili/dispersal. These simulation data were analyzed and visualized using custom Wolfram Mathematica scripts and R scripts (https://github.com/chenxiaolili/dispersal).

## Competing Interests

The authors declare that they have no conflict of interest.

## Acknowledgments

We thank the entire Wu lab for critical discussion of our manuscript. We wish to thank Dr. T. Juelich (UCAS, Beijing) for linguistic assistance during the preparation of this manuscript. This work was supported by the National Key R&D Program of China (2018YFA0902100 and 2021YFA0910300), the National Natural Science Foundation of China (91951204, 32130004, 32161133023, and 32170113), Sino Swiss Science and Technology Cooperation (SSSTC) Program (IZLCZ0_206044), and the High-performance Computing Platform of Peking University.

## Notes

### Competing Interest Statement

The authors have declared no competing interest.

### Summary of Updates

More results

