## Supplemental file for "High immigration rates critical for establishing emigration-driven diversity in microbial communities"

### S1 Construction of community structure detection method

#### S1.1 Validation of DNA extraction method

The 20-strain collection is phylogenetically diverse and spans 20 species across 12 genera and eight families. Since the selected 20 strains are from different genera (Fig. S1), there is a preference for DNA extraction. The cloning library experiment was performed to preliminarily judge the difference in DNA extraction efficiency of the 20 strains. After 20 strains were mixed in the same proportion, total DNA was extracted, and the V3V4 region sequence of 16S rRNA was then amplified. The amplified sequence was constructed into 4 parallel clone libraries. Clone library 1 contained species Ag, Ac, Av, Cc Sf, and Lm. Clone library 2 was consisted of species Ba, Av, Ca, Ka, Fb, and Pk. Species Bp, Av, Fg, Kr, Pa, and Ki were included in clone library 3. Clone library 4 was composed of species Ao, At, Av, Lf, and Pg. A total of 60 single clones were picked and sequenced to determine the actual number of clones of different strains in each clone library. The results showed that different strains differed in the number of clones in the clone library, which proved that there were indeed differences among 20 strains in the DNA extraction efficiency (Figure S2).

#### S1.2 DNA extraction efficiency correction

To accurately determine the DNA extraction efficiency of 20 strains for later community composition correction, reference strain *Agrobacterium tumefaciens* GV 3101 was introduced. According to the preliminary differences in the DNA extraction efficiency of the 20 strains in the clone library results (Figure S2), the 20 strains were divided into two groups (Table S2), the group with higher DNA extraction efficiency and the group with lower DNA extraction efficiency, respectively. Frozen stocks of 20 trains and reference strain *A. tumefaciens* GV

3101 were streaked out on LB agar Petri dishes, grown at 30℃ until strains grew up. Single colonies were picked and each strain was grown separately in 3 ml LB broth until their stationary phases, and then 2% inoculum was transferred to 20 ml LB broth for 24 h. Cells were then collected by centrifuging at 3000 g for 10 minutes and were washed three times using M9 minimal medium without carbon source. The individual cultures were then adjusted to an $OD_{600}$ = 0.5. Each strain was mixed with *A. tumefaciens* GV 3101 according to the relative proportion gradient in Table S2. The DNA of the mixtures was extracted. V3-V4 sequence region was then amplified to obtain the relative abundance of *A. tumefaciens* GV 3101 and the target strain at each concentration. A linear fit between the ratio obtained by sequencing and the initial mixing ratio was performed. The slope of the fit was defined as the extraction correction coefficient of the target strain (Table S2).

### S2 Supporting Figures

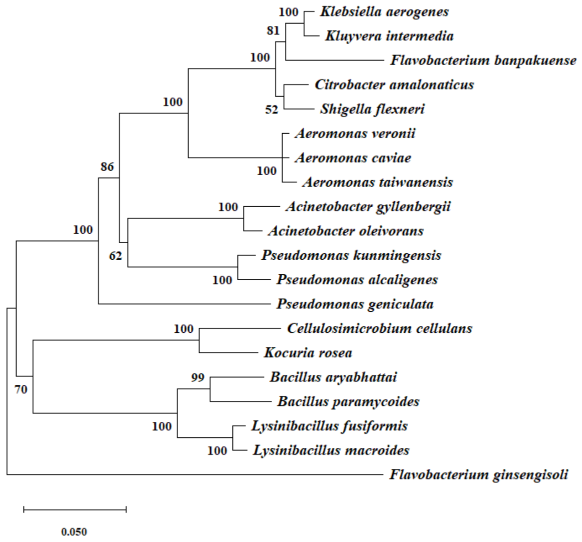


**Figure S1** Phylogenetic tree of the 20 bacterial species used in this study. The tree was constructed based on the 16S rRNA genes and the branch lengths indicated the number of substitutions per base pair. Scores on nodes indicate the posterior probability.

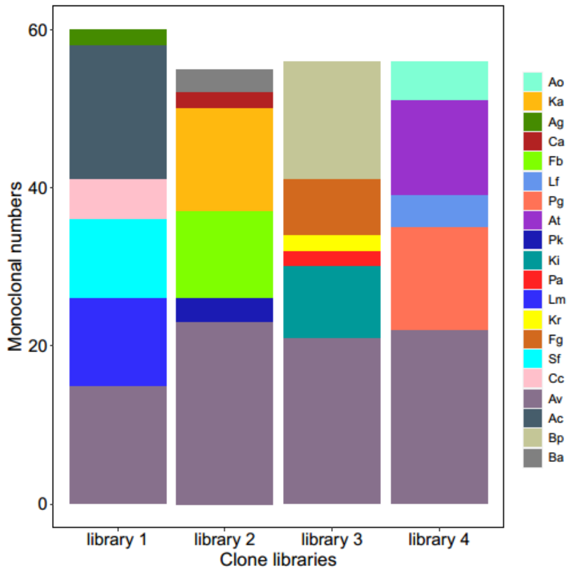


**Figure S2** Statistics of monoclonal numbers in the clonal library. After 20 strains were mixed in the same proportion, total DNA was extracted, and then the sequence of the V3V4 region of 16S rRNA was amplified. The clone libraries were constructed using amplified sequences. About 60 single clones were picked from each clone library and sequenced to obtain the actual number of clones of different strains in the clone library. The cloning library experiments preliminarily showed the difference in DNA extraction efficiency of 20 strains.

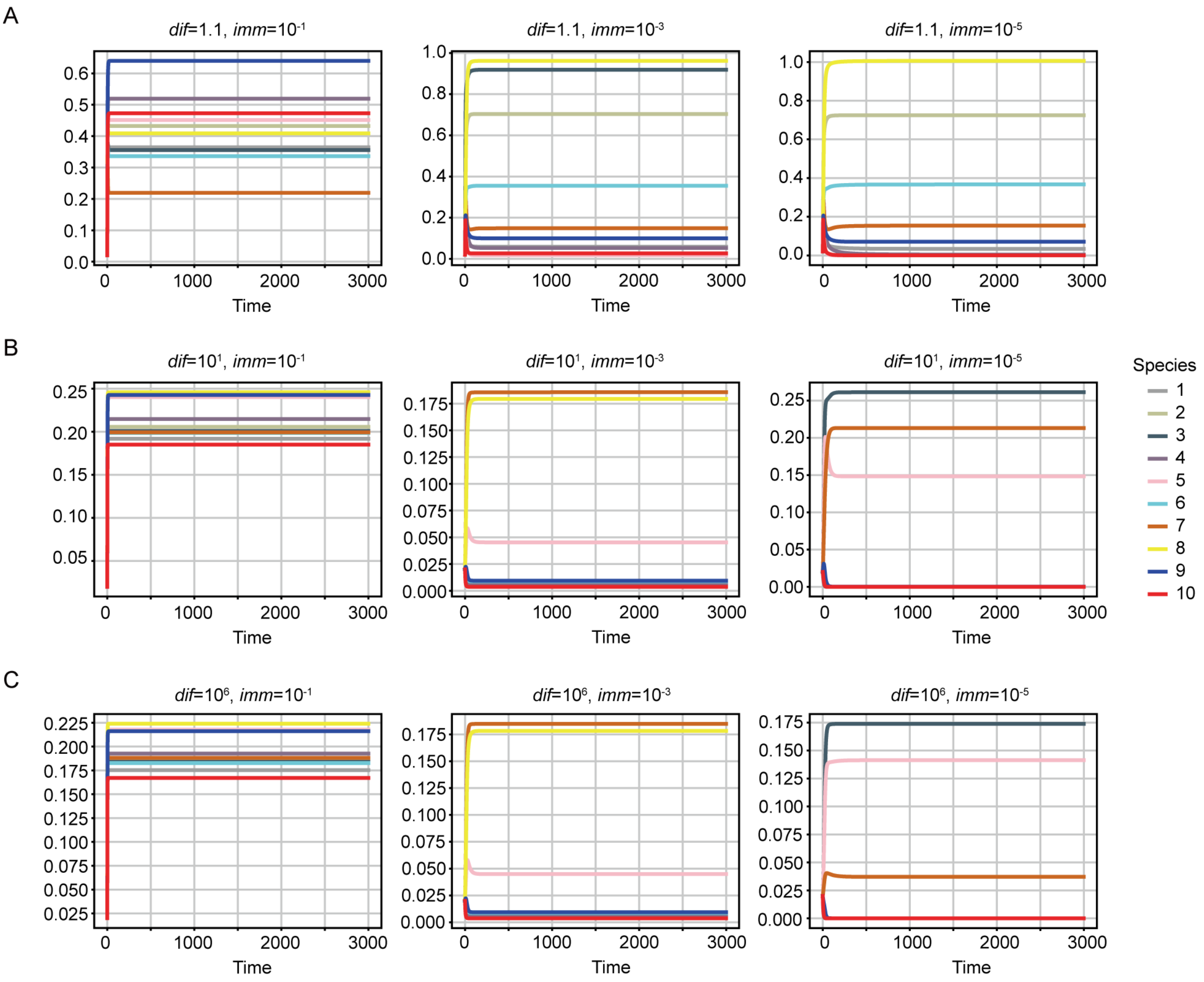


**Figure S3** The dynamics of the microbial communities in simulations under a range of dilution factors (1.1 - $10^6$, three rows) and immigration rates ($10^{-5}$ – $10^{-1}$, three columns). We set the time to $3\times10^3$ for each simulation, which allowed the microbial community to reach equilibrium.

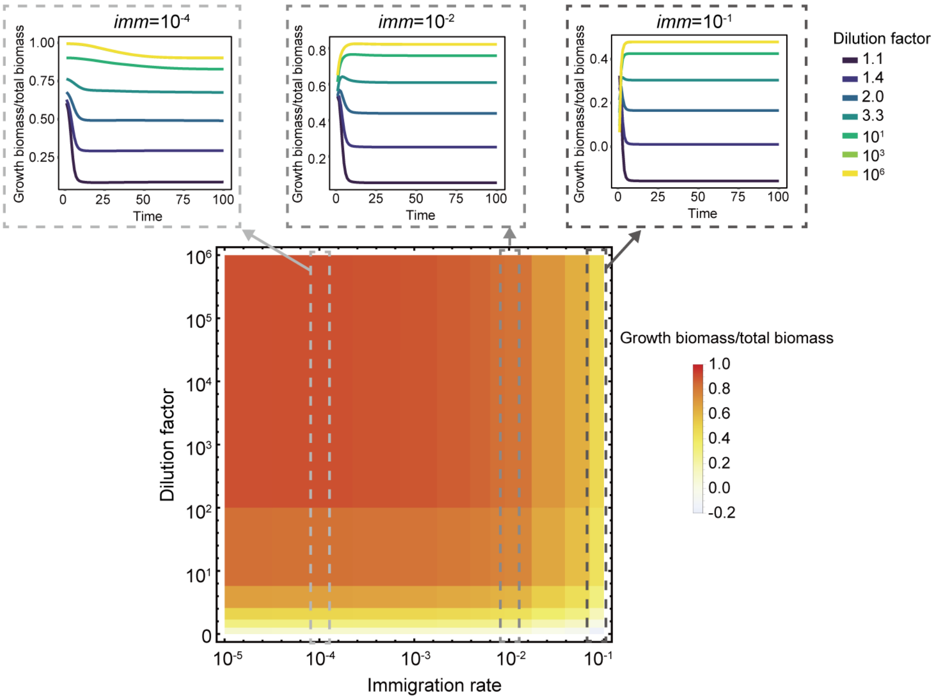


**Figure S4** Effects of immigration rates and dilution factors on the fraction of growth biomass during community succession. The total biomass of a community in Equation [1] is composed of the growth biomass ($N_i\, r_i(1-\frac{N_i}{K_i}-\frac{\sum_j c_{ij}N_j}{K_i})$) and the dispersal biomass ($-\delta N_i + imm$). The three figures in top row represent the dynamics of growth of the community members in the low, middle, and high immigration rates (from left to right). The y-axis is obtained through dividing the growth biomass by the total biomass. As shown in the heatmap, when the dilution factor was more than $10^1$, the fractions of the community growth were more than 0.39 across all immigration rates ($10^{-5}$ – $10^{-1}$), which demonstrated that the term in parentheses ($N_i\, r_i(1-\frac{N_i}{K_i}-\frac{\sum_j c_{ij}N_j}{K_i})$) played an important role during community succession.

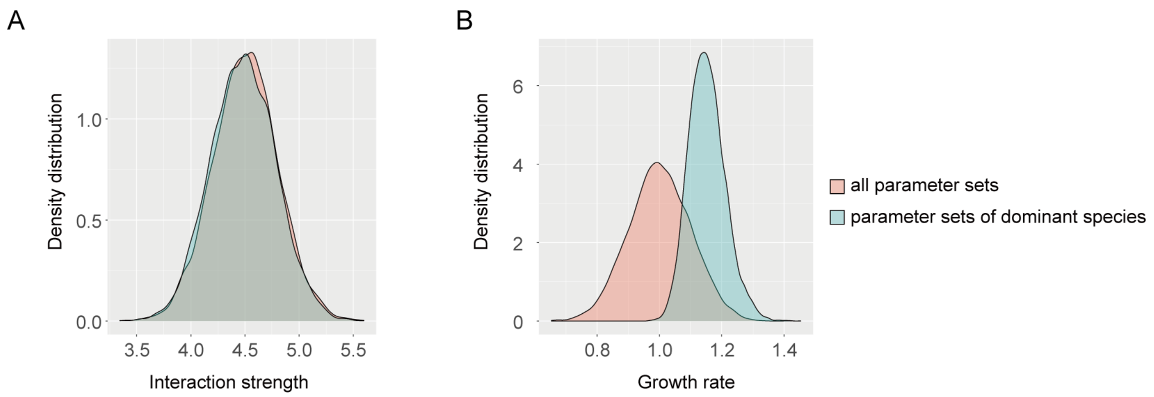


**Figure S5** Parameter settings of interaction strength (A) and growth rate (B) in a total of 10 species and dominant species, respectively. Here, the dominant bacteria had no advantage in interaction strength but had a growth rate advantage. The interaction strength of dominant bacteria was obtained by calculating the sum of the interaction strength of other species on advantaged bacteria in a community.

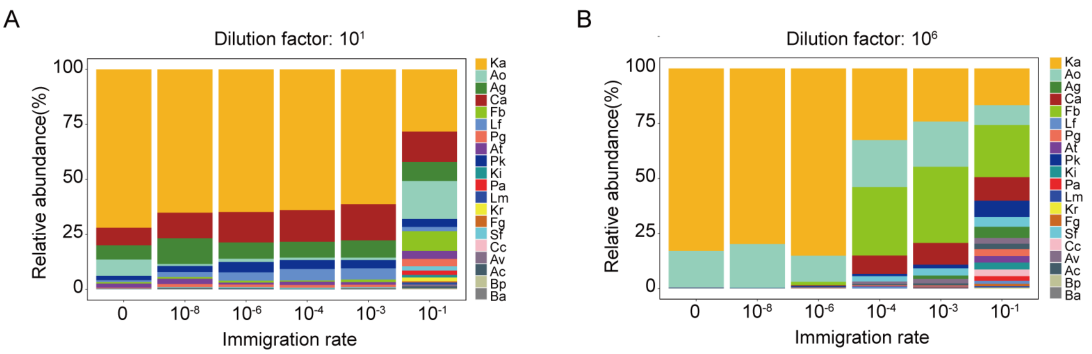


**Figure S6** Effects of immigration rates on the diversity of bacterial communities. The community composition with dilution factors of $10^1$ (A) and $10^6$ (B), respectively.

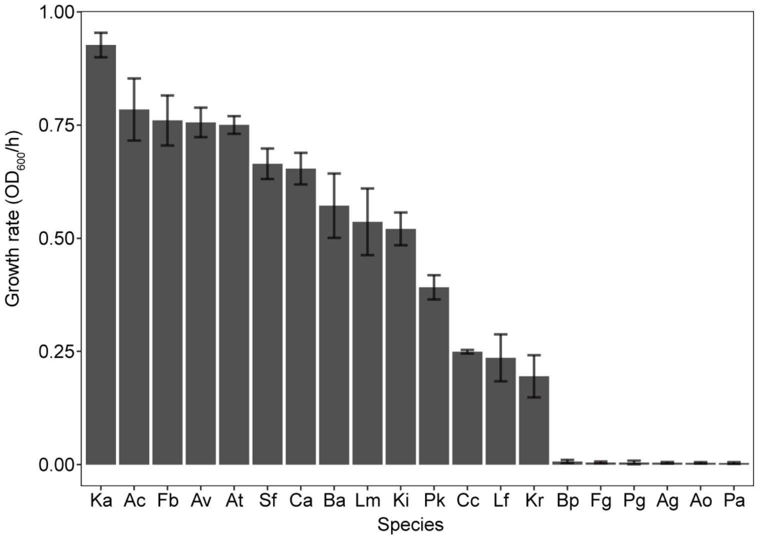


**Figure S7** Experimentally measured growth rates of 20 strains. They display a variety of growth rates, which reflects the growth diversity of natural microbial communities.

**S3 Supplementary Tables**

**Table S1** Identities of 20 strains in our experiments

| Number | The most similar strain | Name for short |
| --- | --- | --- |
| 1 | *Acinetobacter gyllenbergii* CIP 110306 | Ag |
| 2 | *Acinetobacter oleivorans* DR1 | Ao |
| 3 | *Bacillus aryabhattai* B8W22 | Ba |
| 4 | *Bacillus paramycoides* NH24A2 | Bp |
| 5 | *Aeromonas caviae* CECT 838 | Ac |
| 6 | *Aeromonas taiwanensis* LMG 24683 | At |
| 7 | *Aeromonas veronii* CECT 4257 | Av |
| 8 | *Cellulosimicrobium cellulans* LMG 16121 | Cc |
| 9 | *Citrobacter amalonaticus* CECT 863 | Ca |
| 10 | *Klebsiella aerogenes* KCTC 2190 | Ka |
| 11 | *Shigella flexneri* ATCC 29903 | Sf |
| 12 | *Flavobacterium ginsengisoli* DCY54 | Fg |
| 13 | *Flavobacterium banpakuense* 15F3 | Fb |
| 14 | *Kocuria rosea* DSM 20447 | Kr |
| 15 | *Lysinibacillus fusiformis* NBRC 15717 | Lf |
| 16 | *Lysinibacillus macroides* DSM 54 | Lm |
| 17 | *Pseudomonas kunmingensis* HL22-2 | Pk |
| 18 | *Pseudomonas geniculata* ATCC 19374 | Pg |
| 19 | *Pseudomonas alcaligenes* NBRC 14159 | Pa |
| 20 | *Kluyvera intermedia* NBRC 102594 | Ki |

**Table S2** Strain grouping in the determination of DNA extraction efficiency

| Group | Species | Strain tested/ *A. tumefaciens* GV3101 |
|---|---|---|
| Species with higher extraction efficiency | Ag, Ao, Ba, Cc, Ca, Kr, Lf, Pk, Pa | 1:10, 1:5, 1:2, 1:1, 2:1, 5:1 |
| Species with lower extraction efficiency | Bp, Ac, At, Av, Ka, Sf, Fg, Fb, Lm, Pg, Ki | 1:1, 2:1, 5:1, 10:1, 15:1, 30:1 |

**Table S2** Correction formulas for DNA extraction efficiencies of 20 species.

| No. | Species | Formula | $R^2$ |
|---|---|---|---|
| 1 | Ag | y=1.73+1.92x | 0.73 |
| 2 | Ao | y=-2.02+2.66x | 0.96 |
| 3 | Ba | y=-4.78+42.5x | 0.96 |
| 4 | Bp | y=-0.3+2.36x | 0.99 |
| 5 | Ac | y=-0.485+1.25x | 0.91 |
| 6 | At | y=-0.621+1.34x | 0.88 |
| 7 | Av | y=-0.446+0.839x | 0.96 |
| 8 | Cc | y=-0.811+63.1x | 0.96 |
| 9 | Ca | y=-1.1+1.96x | 0.94 |
| 10 | Ka | y=-0.48+1.64x | 0.93 |
| 11 | Sf | y=-0.39+1.19x | 0.97 |
| 12 | Fg | y=-0.373+1.51x | 0.98 |
| 13 | Fb | y=-0.666+1.81x | 0.95 |
| 14 | Kr | y=-0.208+48.4x | 0.96 |
| 15 | Lf | y=-1.66+7.04x | 0.98 |
| 16 | Lm | y=-0.479+1.46x | 0.95 |
| 17 | Pk | y=0.429+1.05x | 0.96 |
| 18 | Pg | y=-0.707+1.67x | 0.95 |
| 19 | Pa | y=-2.32+1.5x | 0.95 |
| 20 | Ki | y=-0.315+1.51x | 0.94 |
